# Perspective: Genomic inference using diffusion models and the allele frequency spectrum

**DOI:** 10.1101/375048

**Authors:** Aaron P. Ragsdale, Claudia Moreau, Simon Gravel

## Abstract

Evolutionary, biological, and demographic processes combine to shape the variation observed in populations. Understanding how these processes are expected to influence variation allows us to infer past demographic events and the nature of selection in human populations. Forward models such as the diffusion approximation provide a powerful tool for analyzing the distribution of allele frequencies in contemporary populations due to their computational tractability and model flexibility. Here, we discuss recent computational developments and their application to reconstructing human demographic history and patterns of selection at new mutations. We also reexamine how some classical assumptions that are still commonly used in inference studies fare when applied to modern data. We use whole-genome sequence data for 797 French Canadian individuals to examine the neutrality of synonymous sites. We find that selection can lead to strong biases in the inferred demography, mutation rate, and distributions of fitness effects. We use these distributions of fitness effects together with demographic and phenotype-fitness models to predict the relationship between effect size and allele frequency, and contrast those predictions to commonly used models in statistical genetics. Thus the simple evolutionary models investigated by Kimura and Ohta still provide important insight into modern genetic research.

---

The scope of human sequencing projects has grown rapidly in recent years to include hundreds of thousands of samples from populations around the globe. Analyzing such data requires tools that can handle large numbers of samples, take into account a wide range of demographic and evolutionary scenarios, and make predictions about relevant statistical properties of the data. In this review we focus on the distribution of frequencies of derived alleles in a population (also known as the allele frequency spectrum or AFS). The simplicity of the statistic, coupled with its sensitivity to demographic and evolutionary processes, has led to its wide use in genomic inferences, including past population demography (Williamson et al., 2005; Gutenkunst et al., 2009), the distribution of fitness effects for new mutations (Boyko et al., 2008; Kim et al., 2017; Huber et al., 2017), and expected patterns of diversity in spatially expanding populations and tumors (Peischl and Excoffier, 2015; Fusco et al., 2016).

Given an evolutionary model, the expected AFS can be estimated through a diversity of methods. Individual-level, population-wide simulation is a conceptually straightforward choice, but until recently simulations were limited to very short subsets of the genome or to very small populations.

Diffusion approaches were introduced as an analytically and computationally tractable approach to model the distribution of allele frequencies without the need to track individuals or alleles through evolution. Following early work by Kolmogorov, Fisher, and Wright, the diffusion equation in the context of population genetics was extensively studied by Kimura (1964), Ohta and Kimura (1971)and contemporaries. Many classical results of population genetics trace their roots to the diffusion approaches, including the expected distribution of allele frequencies in a population, the probability and time to fixation of new mutations, and the evolution of linkage disequilibrium, e.g. (Kimura and Crow, 1964; Kimura and Ohta, 1969; Hill and Robertson, 1968).

The 1980s brought about the rise of approaches based on coalescent theory, which traces lineages backward in time and computes statistics through the expected branch lengths of the resulting trees. The coalescent gained popularity due to its ease of implementation and speed of simulation, quickly becoming a go-to simulation engine (Hudson, 2002). Recent advances allow for coalescent-based simulations of neutral evolution with many samples, many populations, and large genomes (Kelleher et al., 2016). However, genome-wide coalescent simulations are still limited to a narrow range of selective scenarios.

Rapid improvements in algorithms and computational power over the past two decades led to the resurgence of forward models, which allow for large-scale simulations with various selection regimes. Together with genome-level simulations (Hernandez, 2008; Thornton, 2014; Haller and Messer, 2017), the classical diffusion approximation found new life through numerical approaches to compute the AFS for non-equilibrium multi-population demography, selection and dominance (Gutenkunst et al., 2009; Lukic and Hey, 2012; Jouganous et al., 2017).

Here we discuss inference of demography and selection from the AFS and how these combine to influence our predictions of the architecture of complex traits (e.g. epilepsy, height). We use a large whole-genome sequenced cohort of French Canadians and data from The 1000 Genomes Project Consortium (2015) to examine how assumptions commonly used in genome-wide inference fare when applied to large cohorts. We consider in particular the assumption of neutrality at synonymous sites, its effect on inferences of demography and distributions of fitness effects for new mutations, and the relationship between variant frequency and effect size.

## Inference from the AFS

Diffusion-based forward approaches often focus on modeling the distribution of allele frequencies (AFS). Given a sample of size **n** = (*n*_1_, *n*_2_,…, *n_p_*) haploid copies of the genome in *p* populations, the frequency of the derived allele at a biallelic locus can be described as a vector **i** = (*i*_1_, *i*_2_,…, *i_p_*), where *i_k_* ∈ {0, 1,…, *n_k_*} is the number of derived alleles in population *k.* The AFS Φ_**n**_(**i**) records the number of variants whose derived allele has frequency **i**.

To perform inference given an observed AFS from sequencing data, a straightforward approach is to (1) propose a parameterized model, such as historical demography with size changes, splits and migrations, (2) compute the expected AFS under that model for a given parameter set, (3) compare the expected and observed spectra to obtain the likelihood of those parameters, and repeat (2) and (3) to optimize over model parameters and find the best fit to the data. These parameters typically include mutation rates, population size parameters, migration rates and admixture events, and selection and dominance. The optimization procedure may be carried out through standard minimization algorithms, and a number of inference software include built-in composite-likelihood based inference frameworks (Gutenkunst et al., 2009; Coffman et al., 2016; Jouganous et al., 2017; Kamm et al., 2018). In this process, the expected AFS is often computed for a large number of evolutionary histories, requiring efficient computation of the AFS.

## The diffusion approximation

The classical Wright-Fisher (WF) model describes the forward evolution of a biallelic locus in a population of *N* diploid individuals. The diffusion approach approximates allele frequency dynamics under the discrete WF model by a continuous function *ϕ*(*x*, *t*), the density of loci with allele frequencies *x* ∈ (0, 1) at time *t*. For a population of ancestral size *N* and relative size history 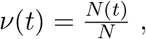, population size-scaled selection coefficient *γ* = 2*Ns*, and dominance coefficient *h*, the evolution of *ϕ* approximately follows the diffusion equation

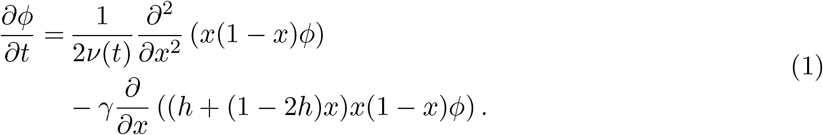

Under the infinite sites assumption (Kimura, 1969), new mutations continually introduce density at frequency 1*/*(2*Nν*(*t*)) with rate proportional to mutation rate *θ* = 4*Nu*.

To compare with data from a finite sample of size *n*, we can then compute the expected *sample* AFS Φ_*n*_(*i*) by integrating *ϕ*(*x, t*) over allele frequencies assuming binomial sampling:

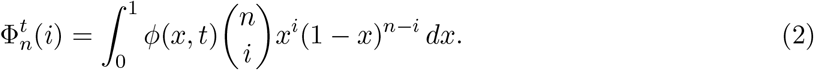

Analytic solutions to Eq. 1 typically exist only for simple, steady-state models. Otherwise, we must turn to numerical solutions. Gutenkunst et al. (2009)proposed to solve Eq. 1 by finite differences, and their popular program *∂*a*∂*i can compute the joint AFS for up to three populations with population size changes, continuous migration, selection and dominance. *∂*a*∂*i also implements likelihood-based optimization package to perform inference on observed data. It is still commonly used to infer demographic histories (e.g., (Hsieh et al., 2016; Mondal et al., 2016)) and patterns of selection for new mutations (e.g., (Ragsdale et al., 2016; Kim et al., 2017; Huber et al., 2017)). Spectral approaches for solving Eq. 1 have also been developed that handle up to four populations (Lukic and Hey, 2012).

The standard diffusion approach and its numerical solution requires two approximations: (1) the approximation of the discrete WF model as a continuous process, and (2) the numerical solution of *ϕ* over a discrete grid (Gutenkunst et al., 2009) or with a truncated sum of polynomials (Lukic and Hey, 2012). Even though the second approximation can in principle be overcome with sufficient computational power, the first approximation can be challenged in very large cohorts (Bhaskar et al., 2014). Evans et al. (2007)proposed a system of ODEs based on the moments of the AFS to avoid the second of these approximations, though their approach leads to numerical instability for large sample sizes. Furthermore, the moment equations only close under neutral evolution – the system of ODEs is infinite under selection. Živković et al. (2015)proposed a truncation approach to close this system of ODEs to simultaneously infer selection strength and recent growth events in a single population with piecewise-constant demography. More recently, Jouganous et al. (2017)proposed a related set of moment equations and moment closure strategy for the frequency spectrum (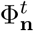) that bypasses both diffusion approximations and allows for computation of the AFS in up to five populations in models with selection and both infinite and finite genome models. This is implemented in the software *moments*, which borrows heavily from *∂*a*∂*i’s user interface and optimization framework.

As is the case for all diffusion-based approaches, going beyond just a handful of populations is difficult, because the number of entries in the frequency spectrum grows exponentially with the number of populations and direct numerical approaches become computationally burdened. For neutral demographic models with more than just a few populations, recent advances to coalescent-based approaches allow for likelihoods to be computed over tens of populations in a tree-shaped demography (i.e. no migration) (Kamm et al., 2017) or up to eight populations with pulse migration events (Kamm et al., 2018) but for fewer samples per population that *∂*a*∂*i or *moments*. In this light, diffusion and coalescent approaches serve complementary roles for simulating and analyzing genomic data, and there are ongoing efforts to improve computational methods for both.

## Evolutionary inference and human history

Understanding the relative roles of demography and selection in shaping present-day diversity is important from anthropological, evolutionary, and biomedical perspectives. To minimize the impact of selection and focus on demography, a well-trodden approach is to focus on putatively neutral sites (Gutenkunst et al., 2009; Gravel et al., 2011; Patterson et al., 2012; Kamm et al., 2018).

One question of particular interest is how modern humans spread across the globe, including the timings of population splits and rates of migration between populations. The popular Out-of-Africa model of human history was proposed by Gutenkunst et al. (2009)and its parameters inferred using *∂*a*∂*i (and further refined as more data has become available (Gravel et al., 2011; Tennessen et al., 2012; Jouganous et al., 2017)). For example, Fig. 2C shows a demographic model inferred from synonymous sites from the 1000 Genomes project data in a panel of world-wide populations (Yoruba from Nigeria (YRI), Utah residents of north-west European ancestry (CEU), Han Chinese from Beijing (CHB), and Japanese from Tokyo (JPT)) byJouganous et al. (2017). In general, populations with non-African ancestry show reduced levels of overall diversity consistent with an Out-of-Africa bottleneck and a relative excess of singletons suggesting recovery and recent growth (Fig. 2D). This contemporary growth has been the subject of much recent investigation (Gazave et al., 2014; Gao and Keinan, 2016), and increasing sample sizes provide insight into the distribution of rare variants, which is particularly sensitive to recent demographic events. It is also sensitive to negative selection which, just as population growth, increases the proportion of rare variants (Fig. 1E).

**Figure 1:**
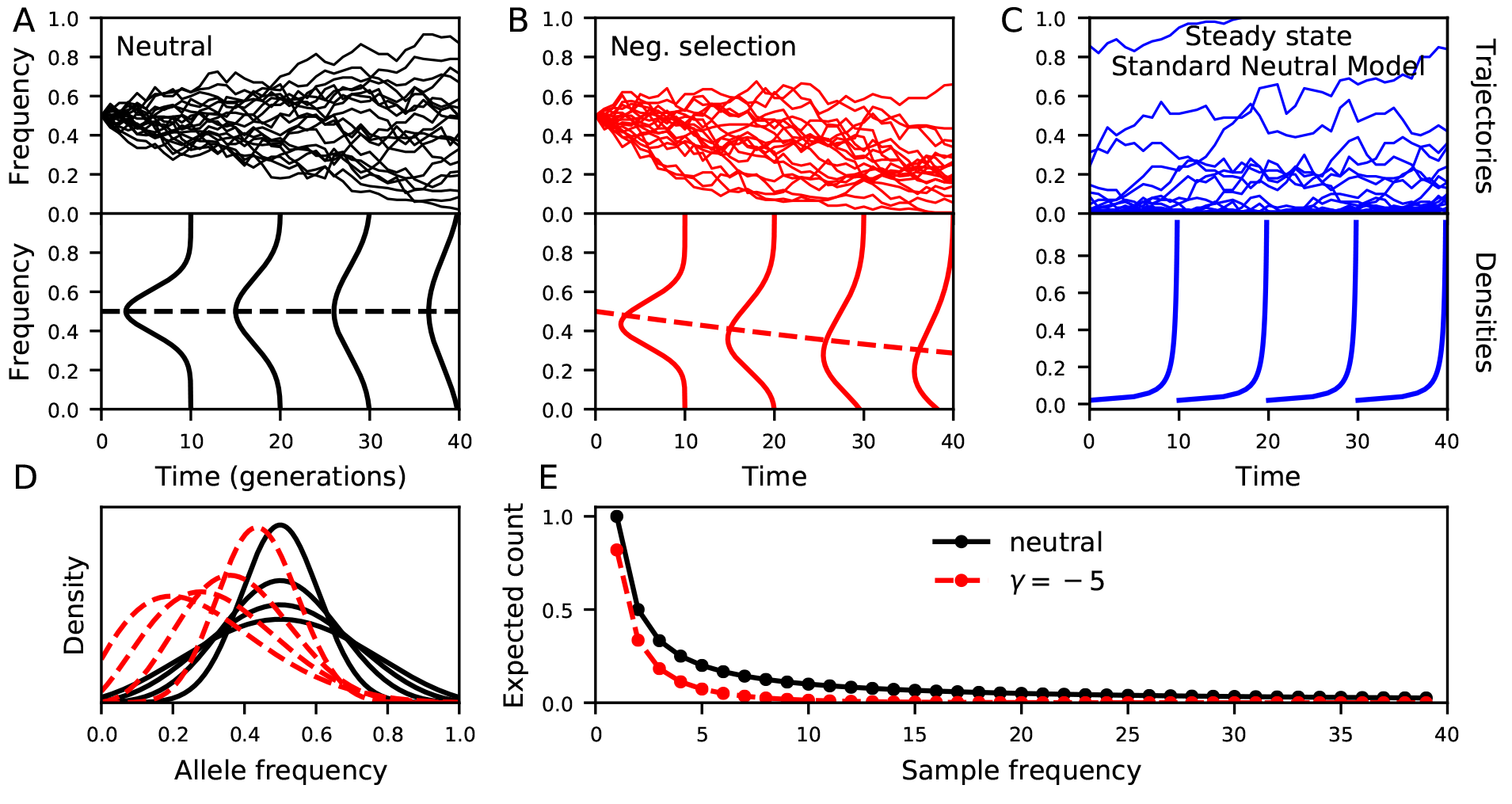
Sample trajectories and the AFS. (A) Neutral mutation trajectories starting from frequency 1*/*2 (above) and the associated diffusion approximation for their frequencies over time (frequency on vertical axis, density on horizontal). While the expected frequency for a neutral mutation does not change over time (equal probability to increase or decrease), variance due to finite population sizes leads to the eventual fixation or loss of the allele. (B) Selected variants tend to increase when positively selected and decrease when negatively selected. The diffusion approximation easily accounts for the effect of selection on the expected distribution of allele frequencies. (C) Under the standard neutral model, mutations give rise to new variants beginning with frequency 1*/*2*N*, which then segregate until they fix or are lost. The continual influx of new mutations and constant size demography leads to a steady state distribution of allele frequencies. (D) The variance of allele frequencies increases over time until the mutation is either fixed or lost in the population. (E) The sample AFS. Negatively selected variants are lost more rapidly and skewed to lower frequency than neutral variants.

**Figure 2:**
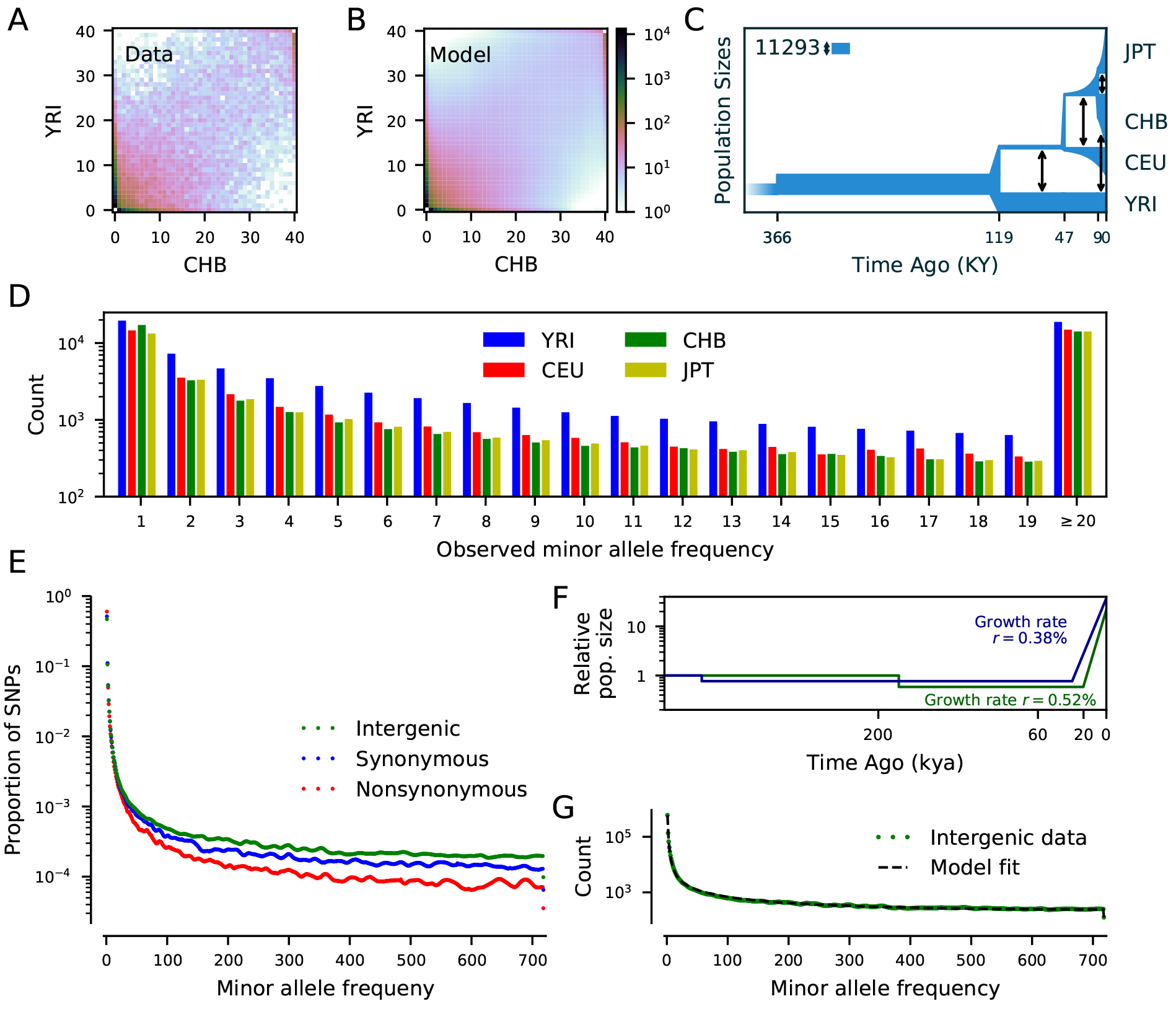
Human demography inferred from the AFS. Comparison of (A) the synonymous joint AFS between Yoruba and Han Chinese samples in the Thousand Genomes Project data and (B) the model as expected under an inferred demographic history (C). (D) Comparison of synonymous AFS for four populations from the Thousand Genomes Project, each with ~ 100 sampled individuals (2*n ≈* 200). Populations that experienced bottlenecks due to the Out-of-Africa range expansion harbor fewer low frequency variants, though alleles with MAF≥ 20 are found at similar levels across populations. Recent exponential growth has lead to a comparable numbers of singletons across populations. (E) AFS from the French Canadian population. The synonymous AFS is skewed to lower frequencies relative to intergenic variants; this is likely caused by at least some synonymous variants being under the effect of direct or linked selection. (F) A simple demographic history for the French Canadian population fit to the intergenic (green) and synonymous (blue) AFS in (E). The two histories are qualitatively similar, but inferred times and growth rates differ. Scaling assumed *N_e_* = 11, 293 and mean generation time of 29 years (Gravel et al., 2011). (G) Comparison of the intergenic AFS and the expected AFS from the best fit demographic model in (F).

### Are synonymous variants a good proxy for neutrality?

Inferences on population growth have largely been constrained to “putatively neutral” synonymous variation because exome capture technology made large datasets of high-quality exome data more readily available than whole-genome data. However, the assumption of neutrality for synonymous mutations may not be tenable, as selection at linked nonsynonymous variants may skew the synonymous AFS (Messer and Petrov, 2013; Ewing and Jensen, 2016; Schrider et al., 2016; Cvijović et al., 2018; Torres et al., 2018), and synonymous mutations may themselves be directly under selection.

One obvious way to reduce selection-induced biases in demographic inferences is to focus on variants that are less affected by selection, such as nonconserved intergenic regions far from genes. For example, Gazave et al. (2014)identified a set of such putatively neutral regions which they sequenced in 500 individuals of European ancestry to high coverage, and they inferred growth rates higher than 3% per generation over the last *∼* 150 generations.

Using high-quality whole-genome sequence data, we are now able to directly compare intergenic and synonymous AFS in a given cohort. We used intergenic regions identified as likely neutrally evolving (Arbiza et al., 2012) (total length *L ≈* 81.2 Mb), as well as autosomal protein coding regions (*L ≈* 33.8 Mb) to compare the AFS for nonconserved intergenic, synonymous, and non-synonymous mutations in 797 French Canadian individuals (Fig. 2E, data details in Appendix). If both intergenic and synonymous mutations are neutral, their frequency spectra should match in proportion. However, the synonymous AFS is notably skewed to rare variants when compared to the intergenic AFS. Because these data come from the same sequenced individuals with comparable coverage, we could conclude that either (1) some synonymous variants are directly under negative selection, which skews the AFS to rare variants (Fig. 1E), or (2) selection at linked sites (e.g. nonsynonymous or regulatory variants) is responsible for the skew of the AFS.

To quantify the impact of these differences on demographic inference, we considered a single population demographic model for the French Canadian population fit to both intergenic and synonymous data. The demography included an extended bottleneck followed by recent exponential growth (Fig. 2F), and we compared the inferred growth rates and times of onset of recent growth. The best fit model from intergenic data had a growth rate of 0.52(*±*0.01)% beginning 20.3(*±*0.3) thousand years ago, while the model inferred from synonymous data had a growth rate of 0.38(*±*0.03)% beginning 30.0(*±*1.3) thousand years ago. Even though both models are arguably in qualitative agreement, biases induced by assumptions of neutrality were much larger than that statistical uncertainties caused by the finite amount of data from the sequenced regions. Inference of human demography would be better served by intergenic data, even if whole genome dataset sizes pale in comparison to exome sequencing datasets.

The total number of variants in the AFS depends on the constant *θL*, where *L* is the total length of sequence used to compute the AFS and *θ* = 4*N_e_μ*. If *N_e_* can be estimated from external sources, AFS inference can be used to provide estimates of the mutation rate (Gravel et al., 2013; Kamm et al., 2018). The coding mutation rate inferred from synonymous data, *θ*_cds_ = 0.000720 *±* 0.000025, was far smaller than the rate inferred from intergenic data, *θ*_int_ = 0.00111 *±* 0.00001, so that *μ*_int_*/μ*_cds_ = 1.53. Using the same value of *N_e_* = 11, 293 as above, the per-base per-generation mutation rates would be *μ*_cds_ = 1.59 *×* 10^*−*8^ and *μ*_int_ = 2.45 *×* 10^*−*8^, suggesting that direct or linked selection can lead to substantial underestimation of the coding mutation rate.

## Selection in human populations

### The distribution of fitness effects for new mutations

A fundamental biological parameter determining the amount of deleterious variation in a population is the distribution of fitness effects (DFE) of new mutations. Whereas some model organisms allow for the determination of the DFE through mutagenesis and competition experiments (Firnberg et al., 2014; Wrenbeck et al., 2017), this is impractical (and unethical) for many organisms. We must turn to statistical approaches to infer fitness effects, but this requires the joint modeling of demography, fitness and dominance coefficients. A typical approach first controls for drift by fitting a demographic model to putatively neutral sites, and then fits a model including both drift and a parameterized DFE to putatively functional variants (Boyko et al., 2008; Ragsdale et al., 2016; Kim et al., 2017). Because exon data is readily available, the demographic model is often inferred from synonymous variation, and the DFE inferred from nonsynonymous variation.

### The distribution of fitness effects and the synonymous AFS

We observed above that in a sample of 797 French Canadian individuals, the AFS for synonymous variants is skewed to low frequencies when compared to putatively neutral mutations from intergenic regions. This skew is likely caused by a combination of background selection from linked nonsynonymous or regulatory sites (Ewing and Jensen, 2016; Cvijović et al., 2018) and selection acting directly on synonymous variants. Here we’ll consider the two extreme scenarios (background selection only and direct selection only), and discuss consequences for the distribution of fitness effects, since this will provide us with a range of plausible DFEs.

In the background selection model, synonymous variation is assumed to be purely neutral so the DFE is concentrated at 0. In the direct selection scenario, we can characterize the effect of selection on synonymous variation by fitting a DFE for synonymous mutations by controlling for demography with the intergenic AFS (which we assume to be neutral). Assuming the same mutation rate in intergenic and coding regions and using the Kim et al. (2017)estimate for the ratio *μ*_non_*/μ*_syn_ = 2.31, we inferred a gamma distributed DFE (Fig. 3(C,E)). The majority of synonymous mutations were inferred to be effectively neutral or only slightly deleterious, but under this assumption around 10% of synonymous mutations have selection coefficients *|s| ≥* 10^*−*3^ with *<* 0.1% having *|s| ≥* 10^*−*2^.

**Figure 3:**
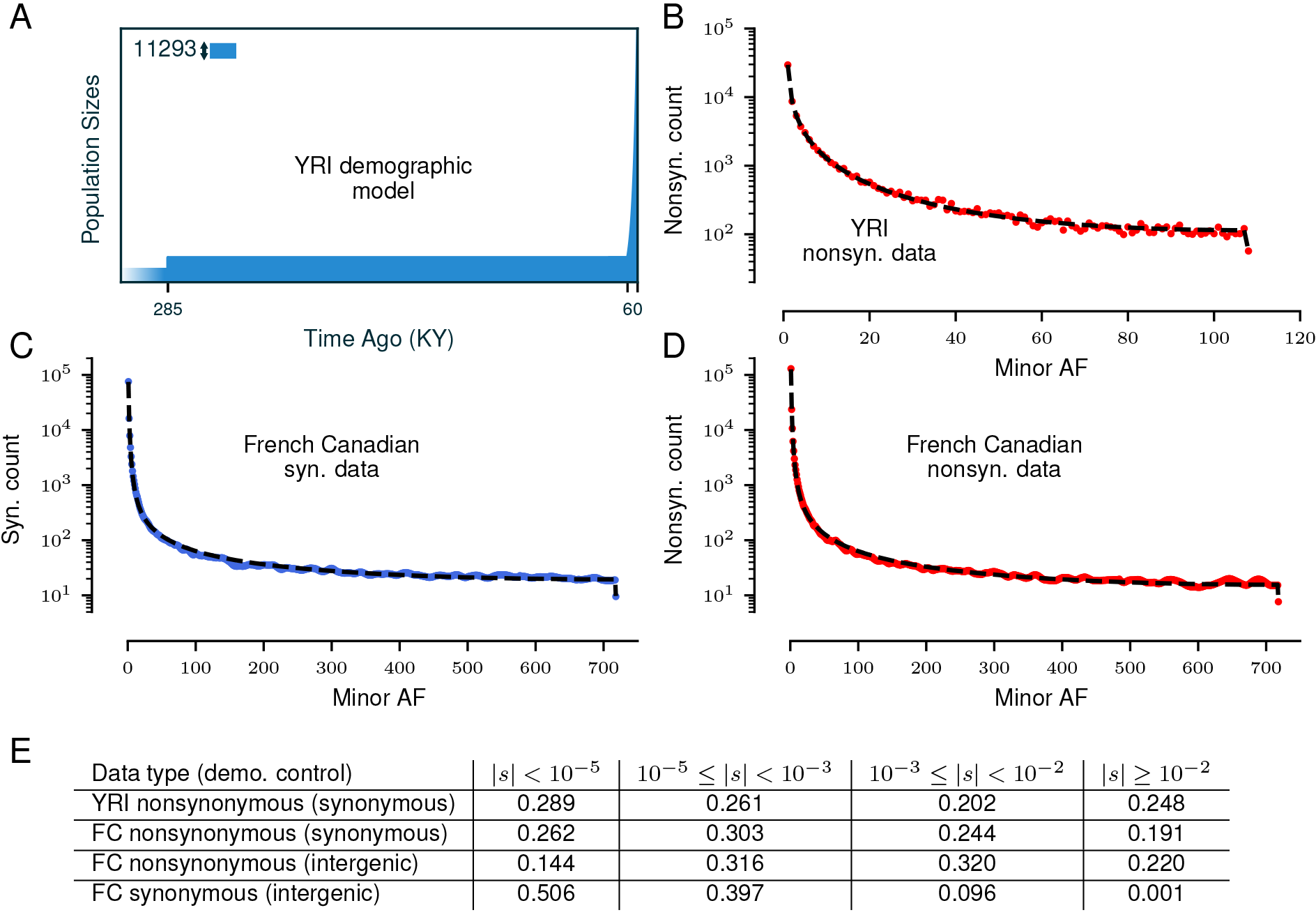
Inference of the DFE for nonsynonymous mutations. Using synonymous and nonsynonymous SNPs from 108 Phase 3 Thousand Genomes Project Yoruba individuals, we (A) fit a single-population demography to the synonymous AFS, and (B) inferred a gamma-distributed DFE (*α* = 0.14, *β* = 2590) to the nonsynonymous AFS, in agreement with Kim et al. (2017). (C) The expected AFS from a gamma-distributed DFE fit to synonymous variants in 797 French Canadian individuals (*α* = 0.14, *β* = 55), after controlling for demography using intergenic loci (Fig. 2F,G). (D) The expected AFS from the DFE fit to nonsynonymous variants in this same population using intergenic control. (E) Binned densities of selection coefficients for new mutations for each inferred DFE. The inference of the DFE for French Canadian nonsynonymous mutations is sensitive to the variants used to control for demography. Intergenic and synonymous variants provide differing fits to the recent growth rate and time of growth onset, and the DFE we infer using intergenic sites as a demographic control (*α* = 0.25, *β* = 699) suggests relatively fewer nonsynonymous mutations are nearly neutral or slightly deleterious while relatively more are moderately to strongly deleterious, compared to the DFE inferred using synonymous mutations as control (*α* = 0.17, *β* = 1050).

We can repeat the same analysis for nonsynonymous variation. In the background selection scenario, nonsynonymous variants probably experience the same amount of background selection as synonymous variants. Even though the frequency spectrum of synonymous variants is possibly skewed by background selection, it can still be used as a control if we assume that the demographic model fitted to the synonymous variants captures the effects of both demography and background selection acting on coding regions. This is the most commonly used approach for estimating DFEs for nonsynonymous variants (Ragsdale et al., 2016; Kim et al., 2017). In the direct selection case, we simply infer the DFE on nonsynonymous variants by using the intergenic model as a control (Fig. 3(C,E)).

The nonsynonymous DFEs inferred from both models differ in their expected distributions. The mean *|s|* = 0.014 for the direct selection model vs. *|s|* = 0.0078 in the background selection model, although the proportion of variants under strong selection (*|s| ≥* 10^*−*2^) is reduced from 22% to 19%. Resolving the relative contributions of background and direct selection on synonymous variation will therefore be crucial in inferring the DFE – this will require the inclusion of linkage disequilibrium in evolutionary models.

### Selection, effect sizes, and heritability estimates

Because selection and demography affect the distribution of neutral and deleterious variants, they also shape the architecture of traits, that is, the number, frequency, and effect sizes of mutations that contribute to variance in the trait in a population.

A commonly used model in statistical genetics supposes that a phenotype *Y* can be expressed as a sum *Y* = *Xβ* + *e* of genetic contributions *Xβ* and environmental effects *e*. The SNP effect sizes *β* and *e* are assumed to be drawn from normal distributions, and *X* is the genotype matrix. For mathematical convenience (having to do with variance normalization of *X*), the variance of the normal distribution is often assumed to vary with allele frequency, so that 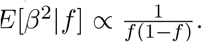. This relationship has the desirable feature that variants with large effect sizes tend to be at low frequency, as expected if functional mutations are deleterious, but its functional form is a rather arbitrary consequence of mathematically convenient assumptions, rather than biological motivation.

These statistical models are used to predict SNP heritability (the proportion of genetic variance that can be explained by variants tagged by a genotyping chip). For example, epilepsy is a highly polygenic trait with many hundreds or potentially thousands of susceptibility loci, each of which typically contributes to just a fraction of a percent to the heritability of the disease (Speed et al., 2014). Using a prior where *E*[*β*^2^*|f*] ∝ 1*/*(*f* (1 *− f*)), Speed et al. (2014)estimated SNP heritability at *∼* 26%, towards the lower range of values suggested by twin studies.

Recent studies have shown that this prior leads to biases in heritability estimates (Speed et al., 2017). This bias may be reduced by binning variants by MAF, so that heritability is estimated independently for each bin (Speed et al., 2017; Evans et al., 2018). While estimates that bin by MAF provide more accurate estimates of heritability for variants across the AFS, they are more computationally demanding and require larger datasets due to the large increase in parameters that must be fit (Evans et al., 2018). For methods that do not stratify by MAF, better estimates could be achieved by including a fudge factor *α* such that *E*[*β*^2^] ∝ (*f* (1 *− f*))^*α*^ (Speed et al., 2017; Zeng et al., 2018). Zeng et al. (2018)estimated *α* to range between *−*0.6 and 0 across a few dozen traits. This indicates the presence of natural selection, but weaker than under the common *α* = *−*1 assumption. Allowing *α* to vary helps reduce the bias compared to fixing *α* = *−*1. But does it correctly account for the relationship between effect size and allele frequency? To answer this question, we need an evolutionary model tying evolution and the distribution of fitness effects, which we discussed above, but also the relationship between variant fitness coefficients and effect size.

If the trait of interest were reproductive fitness, then the distribution of fitness effects would correlate perfectly with the distribution of effect sizes. Using the DFE inferred above for YRI nonsynonymous variation, for example, we find the relationship between effect size and frequency shown in Fig. 4 (*ρ* = 1), which departs notably from (*f* (1 *− f*))^*α*^ models.

**Figure 4:**
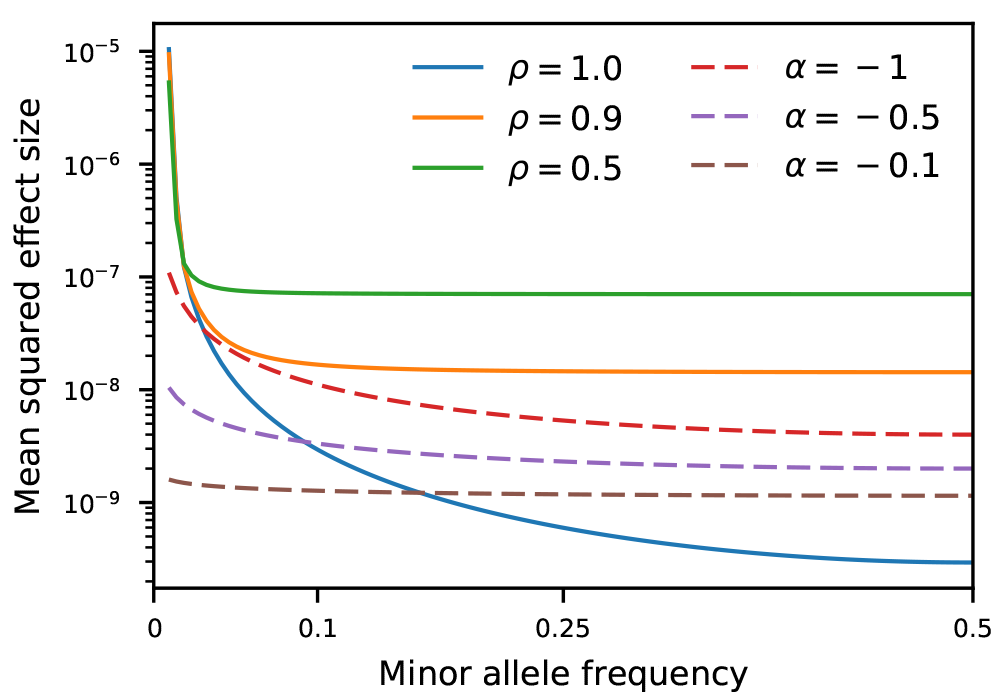
Comparison of population genetic predictions and statistical genetics inferences. Using the DFE inferred above from YRI data, we use the Uricchio et al. (2016)model to predict *β*^2^ as a function of MAF (solid curves) for varying values of *ρ* (larger values of *ρ* imply lower levels of pleiotropy in this model). We compare to the relationship *β*^2^ ∝ (*f* (1 – *f*))^*α*^ (dashed curves), the functional form assumed by Speed et al. (2017)andZeng et al. (2018), for three plausible values of *α*. For these particular models, the predicted relationship from population genetics and assumptions from empirical studies notably differ.

In a complex trait such as epilepsy or height, *s* is unlikely to correlate perfectly with effect size because of pleiotropy or simply because the effect of selection on the trait is absent or nonlinear. Many models have been recently proposed in the hope of tying evolutionary models to GWAS findings (Uricchio et al., 2016; Sanjak et al., 2017; Simons et al., 2018). Using the model from Uricchio et al. (2016), together with our inferred demography and DFE, we find distributions of effect sizes shown as solid lines on Fig. 4. They show a much greater difference in phenotypic effects between common and rare variants but also flatter profiles for common variants compared to (*f* (1 *− f*))^*α*^ models, leading to different predictions about the architecture of complex traits.

One possible innocuous reason for the discrepancy is that we considered two different classes of sites: noncoding variants for the evolutionary model, and genome-wide diversity for the (*f* (1 *− f*))^*α*^ empirical relationship. Yet, at least for nonsynonymous variants, allowing for *α* to vary in the (*f* (1 *− f*))^*α*^ model is not enough to capture the relationship between allele frequency and effect size predicted by the Uricchio et al. (2016)model. Both models feature arbitrary choices that may be completely wrong. Fortunately, we now have both the data and models to directly compare predictions from statistical and evolutionary genetic models, and studies discussed in this section have started to take advantage of them.

## Conclusion: what can we learn from the AFS?

The distribution of allele frequencies is a simple summary statistic for genomic diversity. It is a relatively smooth and monotonous function from which we are trying to extract a lot of information, ranging from population sizes, split times, migration rates, the distribution of fitness effects, and the relationship between trait effect sizes and fitness. Even though the AFS does contain information about all these processes, there are more parameters to learn than data points, and this raises the question of how much meaningful information can be extracted from the data.

This question has received considerable theoretical interest in the simplified context of inferring population sizes from the AFS in a single, neutrally evolving population. An elegant theoretical study by Myers et al. (2008)showed that distinct size change histories could lead to identical AFS, no matter how large the sample size or number of markers studied. This presents a fundamental problem for inference: if multiple demographic histories have the same expected AFS, how can we be certain that our inferred demography has converged to the correct function? Since the Myers et al. (2008)study, there has been back and forth regarding limits to inference when size histories are constrained to biologically realistic models (Bhaskar and Song, 2014), placing bounds on the convergence rates of inference from the AFS (Terhorst and Song, 2015), and limits to inference with realistic data from a genome of finite length (Rosen et al., 2017; Baharian and Gravel, 2018). As may be expected, ancient and pre-bottleneck histories are difficult to accurately reconstruct from the AFS or other summaries of the data because of the lack of contemporary diversity originating from those epochs, and recent historical events are more tightly constrained by the data.

Similarly, any inference problem using genomic data is bound to have large regions of uncertainty. If the space of allowed models is sufficiently large, parameter uncertainty will reveal uncertain features of the model. However, it is easy to accidentally parameterize a model in a way that masks uncertainties. Assuming neutrality, or smoothness of population histories, or a particular relationship between fitness and phenotype effects can forbid plenty of plausible evolutionary scenarios. Yet additional model parameters come at considerable computation and interpretation cost.

Of course, we don’t have a general solution to this conundrum, but there is a lot that we can learn by building simple evolutionary models that describe the data well. Only by building layers of simple models for demography, selection, and variant effect sizes were we able to compare results coming from evolutionary and statistical genetics (and find that they disagree!). Thus old-school diffusion-based approaches, taken together with forward simulations, coalescent approaches, and empirical work on large cohorts, still have a lot to tell us about human genomic diversity.

## Acknowledgements

We thank Patrick Cossette, Jacques Michaud, Berge Minassian and Simon Girard for sharing allele frequency spectrum data for the French Canadian population. This research was undertaken, in part, thanks to funding from the Canada Research Chairs program.

## Appendix Data

We extracted synonymous variants from the Phase 3 (2013-05-03) thousand genome annotation VCFs, keeping only biallelic SNPs, for the YRI, CEU, CHB and JPT populations. If a SNP had multiple annotations for different transcripts, we considered it synonymous only if every annotation was synonymous. We similarly pulled nonsynonymous SNPs for the YRI individuals from this same dataset, keeping any variant with a nonsynonymous annotations. Single population AFS were built for each population, and we used the synonymous variants to build the four populations joint AFS (Fig. 2).

797 French Canadian individuals from Quebec were sequenced to a minimum 30x coverage genome wide. These individuals were combined cohorts of 400 epilepsy patients (198 are deposited with the accession code EGAS00001002825 (Monlong et al., 2018) and the rest are as of yet unpublished) and 397 unrelated controls sequenced by the same center and platform as part of the Canadian Epilepsy Network (CENet). We filtered out regions from the Thousand Genomes Project strict mask and sites that were significantly out of Hardy-Weinberg equilibrium (*p <* 0.001, *χ*^2^ test). We used intergenic regions identified by the neutral region explorer (http://nre.cb.bscb.cornell.edu/nre/) (Arbiza et al., 2012), keeping variants at least 0.2 cM from known coding regions and regulatory elements and with background selection coefficient *B >* 0.95.

To compare mutation rates, we used the inferred *θ* from demographic fits to both the intergenic and synonymous AFS. For intergenic sites, we used the total length *L* = 81, 209, 832 to find the per-base population-size scaled mutation rate. For synonymous sites, we used the total length of coding regions included in the analysis (*L*_cds_ = 33, 774, 350), accounting for alternative transcripts and overlapping genes. We scaled *L*_cds_ by the ratio of expected synonymous to nonsynonymous mutations across coding regions to obtain the effective *L*_syn_, *μ*_non_*/μ*_syn_ = 2.31 (Kim et al., 2017). This gives *L*_syn_ = 10, 328, 617, which we used to estimate the coding mutation rate from synonymous data.

We used *moments* to fit demographic models to the full four population AFS and the single population YRI and FRC frequency spectra. *Moments* was also used to generate the frequency spectra used to fit DFEs to the YRI and FRC AFS. Confidence intervals were computed using a parametric bootstrap from 1,000 resampled AFS and found using the Godambe Information Matrix (Coffman et al., 2016).

## References

Arbiza, L., Zhong, E., and Keinan, A. (2012). NRE: a tool for exploring neutral loci in the human genome. BMC Bioinformatics, 13(1):301.

Baharian, S. and Gravel, S. (2018). On the decidability of population size histories from finite allele frequency spectra. Theoretical Population Biology, 120(1):42–51.

Bhaskar, A., Clark, A. G., and Song, Y. S. (2014). Distortion of genealogical properties when the sample is very large. Proceedings of the National Academy of Sciences, 111(6):2385–2390.

Bhaskar, A. and Song, Y. S. (2014). Descartes’ rule of signs and the identifiability of population demographic models from genomic variation data. Annals of Statistics, 42(6):2469–2493.

Boyko, A. R., Williamson, S. H., Indap, A. R., Degenhardt, J. D., Hernandez, R. D., Lohmueller, K. E., Adams, M. D., Schmidt, S., Sninsky, J. J., Sunyaev, S. R., White, T. J., Nielsen, R., Clark, A. G., and Bustamante, C. D. (2008). Assessing the Evolutionary Impact of Amino Acid Mutations in the Human Genome. PLoS Genetics, 4(5):e1000083.

Coffman, A. J., Hsieh, P. H., Gravel, S., and Gutenkunst, R. N. (2016). Computationally Efficient Composite Likelihood Statistics for Demographic Inference. Molecular Biology and Evolution, 33(2):591–593.

Cvijović, I., Good, B. H., and Desai, M. M. (2018). The Effect of Strong Purifying Selection on Genetic Diversity. Genetics, pages 1–52.

Evans, L. M., Tahmasbi, R., Vrieze, S. I., Abecasis, G. R., Das, S., Gazal, S., Bjelland, D. W., de Candia, T. R., Goddard, M. E., Neale, B. M., Yang, J., Visscher, P. M., and Keller, M. C. (2018). Comparison of methods that use whole genome data to estimate the heritability and genetic architecture of complex traits. Nature Genetics, 50(5):737–745.

Evans, S. N., Shvets, Y., and Slatkin, M. (2007). Non-equilibrium theory of the allele frequency spectrum. Theoretical Population Biology, 71(1):109–119.

Ewing, G. B. and Jensen, J. D. (2016). The consequences of not accounting for background selection in demographic inference. Molecular Ecology, 25(1):135–141.

Firnberg, E., Labonte, J. W., Gray, J. J., and Ostermeier, M. (2014). A comprehensive, high-resolution map of a Gene’s fitness landscape. Molecular Biology and Evolution, 31(6):1581–1592.

Fusco, D., Gralka, M., Kayser, J., Anderson, A., and Hallatschek, O. (2016). Excess of mutational jackpot events in expanding populations revealed by spatial Luria-Delbrück experiments. Nature Communications, 7:1–9.

Gao, F. and Keinan, A. (2016). Inference of Super-exponential Human Population Growth via Efficient Computation of the Site Frequency Spectrum for Generalized Models. Genetics, 202(1):235–245.

Gazave, E., Ma, L., Chang, D., Coventry, A., Gao, F., Muzny, D., Boerwinkle, E., Gibbs, R. A., Sing, C. F., Clark, A. G., and Keinan, A. (2014). Neutral genomic regions refine models of recent rapid human population growth. Proceedings of the National Academy of Sciences, 111(2):757–762.

Gravel, S., Henn, B. M., Gutenkunst, R. N., Indap, A. R., Marth, G. T., Clark, A. G., Yu, F., Gibbs, R. A., The 1000 Genomes Project, and Bustamante, C. D. (2011). Demographic history and rare allele sharing among human populations. Proceedings of the National Academy of Sciences, 108(29):11983–11988.

Gravel, S., Zakharia, F., Moreno-Estrada, A., Byrnes, J. K., Muzzio, M., Rodriguez-Flores, J. L., Kenny, E. E., Gignoux, C. R., Maples, B. K., Guiblet, W., Dutil, J., Via, M., Sandoval, K., Bedoya, G., Oleksyk, T. K., Ruiz-Linares, A., Burchard, E. G., Martinez-Cruzado, J. C., and Bustamante, C. D. (2013). Reconstructing Native American Migrations from Whole-Genome and Whole-Exome Data. PLoS Genetics, 9(12).

Gutenkunst, R. N., Hernandez, R. D., Williamson, S. H., and Bustamante, C. D. (2009). Inferring the Joint Demographic History of Multiple Populations from Multidimensional SNP Frequency Data. PLoS Genetics, 5(10):e1000695.

Haller, B. C. and Messer, P. W. (2017). SLiM 2: Flexible, Interactive Forward Genetic Simulations. Molecular Biology and Evolution, 34(1):230–240.

Hernandez, R. D. (2008). A flexible forward simulator for populations subject to selection and demography. Bioinformatics, 24(23):2786–2787.

Hill, W. G. and Robertson, A. (1968). Linkage disequilibrium in finite populations. Theoretical and applied genetics, 38(6):226–31.

Hsieh, P. H., Woerner, A. E., Wall, J. D., Lachance, J., Tishkoff, S. A., Gutenkunst, R. N., and Hammer, M. F. (2016). Model-based analyses of whole-genome data reveal a complex evolutionary history involving archaic introgression in Central African Pygmies. Genome Research, 26(3):291–300.

Huber, C. D., Durvasula, A., Hancock, A. M., and Lohmueller, K. E. (2017). Gene expression drives the evolution of dominance. bioRxiv, pages 1–47.

Hudson, R. R. (2002). Generating samples under a Wright-Fisher neutral model of genetic variation. Bioinformatics, 18(2):337–338.

Jouganous, J., Long, W., Ragsdale, A. P., and Gravel, S. (2017). Inferring the Joint Demographic History of Multiple Populations: Beyond the Diffusion Approximation. Genetics, 206(3):1549–1567.

Kamm, J. A., Terhorst, J., Durbin, R., and Song, Y. S. (2018). Efficiently inferring the demographic history of many populations with allele count data. bioRxiv, pages 1–29.

Kamm, J. A., Terhorst, J., and Song, Y. S. (2017). Efficient computation of the joint sample frequency spectra for multiple populations. Journal of Computational and Graphical Statistics, 26(1):182–194.

Kelleher, J., Etheridge, A. M., and McVean, G. (2016). Efficient Coalescent Simulation and Genealogical Analysis for Large Sample Sizes. PLoS Computational Biology, 12(5):1–22.

Kim, B. Y., Huber, C. D., and Lohmueller, K. E. (2017). Inference of the Distribution of Selection Coefficients for New Nonsynonymous Mutations Using Large Samples. Genetics, 206(1):345–361.

Kimura, M. (1964). Diffusion Models in Population Genetics. Journal of Applied Probability, 1(2):177.

Kimura, M. (1969). The number of heterozygous nucleotide sites maintained in a finite population due to steady flux of mutations. Genetics, 61(4):893–903.

Kimura, M. and Crow, J. F. (1964). The Number of Alleles That Can Be Maintained in a Finite Population. Genetics, 49:725–738.

Kimura, M. and Ohta, T. (1969). The average number of generations until fixation of a mutant gene in a finite population. Genetics, 61(692):763–771.

Lukic, S. and Hey, J. (2012). Demographic Inference Using Spectral Methods on SNP Data, with an Analysis of the Human Out-of-Africa Expansion. Genetics, 192(2):619–639.

Messer, P. W. and Petrov, D. A. (2013). Frequent adaptation and the McDonald-Kreitman test. Proceedings of the National Academy of Sciences, 110(21):8615–8620.

Mondal, M., Casals, F., Xu, T., Dall’Olio, G. M., Pybus, M., Netea, M. G., Comas, D., Laayouni, H., Li, Q., Majumder, P. P., and Bertranpetit, J. (2016). Genomic analysis of Andamanese provides insights into ancient human migration into Asia and adaptation. Nature Genetics, 48(9):1066–1070.

Monlong, J., Girard, S. L., Meloche, C., Cadieux-Dion, M., Andrade, D. M., Lafreniere, R. G., Gravel, M., Spiegelman, D., Dionne-Laporte, A., Boelman, C., Hamdan, F. F., Michaud, J. L., Rouleau, G., Minassian, B. A., Bourque, G., and Cossette, P. (2018). Global characterization of copy number variants in epilepsy patients from whole genome sequencing. PLOS Genetics, 14(4):1–24.

Myers, S., Fefferman, C., and Patterson, N. (2008). Can one learn history from the allelic spectrum? Theoretical Population Biology, 73(3):342–348.

Ohta, T. and Kimura, M. (1971). Linkage disequilibrium between two segregating nucleotide sites under the steady flux of mutations in a finite population. Genetics, 68(4):571–580.

Patterson, N., Moorjani, P., Luo, Y., Mallick, S., Rohland, N., Zhan, Y., Genschoreck, T., Webster, T., and Reich, D. (2012). Ancient admixture in human history. Genetics, 192(3):1065–1093.

Peischl, S. and Excoffier, L. (2015). Expansion load: Recessive mutations and the role of standing genetic variation. Molecular Ecology, 24(9):2084–2094.

Ragsdale, A. P., Coffman, A. J., Hsieh, P., Struck, T. J., and Gutenkunst, R. N. (2016). Triallelic population genomics for inferring correlated fitness effects of same site nonsynonymous mutations. Genetics, 203(1):513–523.

Rosen, Z., Bhaskar, A., Roch, S., and Song, Y. S. (2017). Geometry of the sample frequency spectrum and the perils of demographic inference. bioRxiv, pages 1–21.

Sanjak, J. S., Long, A. D., and Thornton, K. R. (2017). A Model of Compound Heterozygous, Loss-of-Function Alleles Is Broadly Consistent with Observations from Complex-Disease GWAS Datasets. PLoS Genetics, 13(1):1–30.

Schrider, D. R., Shanku, A. G., and Kern, A. D. (2016). Effects of linked selective sweeps on demographic inference and model selection. Genetics, 204(3):1207–1223.

Simons, Y. B., Bullaughey, K., Hudson, R. R., and Sella, G. (2018). A population genetic interpretation of GWAS findings for human quantitative traits. PLOS Biology, 16(3):e2002985.

Speed, D., Cai, N., Johnson, M. R., Nejentsev, S., and Balding, D. J. (2017). Reevaluation of SNP heritability in complex human traits. Nature Genetics, 49(7):986–992.

Speed, D., O’Brien, T. J., Palotie, A., Shkura, K., Marson, A. G., Balding, D. J., and Johnson, M. R. (2014). Describing the genetic architecture of epilepsy through heritability analysis. Brain, 137(10):2680–2689.

Tennessen, J. A., Bigham, A. W., O’Connor, T. D., Fu, W., Kenny, E. E., Gravel, S., McGee, S., Do, R., Liu, X., Jun, G., Kang, H. M., Jordan, D., Leal, S. M., Gabriel, S., Rieder, M. J., Abecasis, G., Altshuler, D., Nickerson, D. A., Boerwinkle, E., Sunyaev, S., Bustamante, C. D., Bamshad, M. J., Akey, J. M., Broad, G. O., Seattle, G. O., and Project, o. b. o. t. N. E. S. (2012). Evolution and Functional Impact of Rare Coding Variation from Deep Sequencing of Human Exomes. Science, 337(6090):64–69.

Terhorst, J. and Song, Y. S. (2015). Fundamental limits on the accuracy of demographic inference based on the sample frequency spectrum. Proceedings of the National Academy of Sciences, 112(25):7677–7682.

The 1000 Genomes Project Consortium (2015). A global reference for human genetic variation. Nature, 526:68–74.

Thornton, K. R. (2014). A C++ Template Library for Efficient Forward-Time Population Genetic Simulation of Large Populations. Genetics, 198(1):157–166.

Torres, R., Szpiech, Z. A., and Hernandez, R. D. (2018). Human demographic history has amplified the effects of background selection across the genome. bioRxiv, pages 1–57.

Uricchio, L. H., Zaitlen, N. A., Ye, C. J., Witte, J. S., and Hernandez, R. D. (2016). Selection and explosive growth alter genetic architecture and hamper the detection of causal rare variants. Genome Research, 26(7):863–873.

Williamson, S. H., Hernandez, R., Fledel-Alon, a., Zhu, L., Nielsen, R., and Bustamante, C. D. (2005). Simultaneous inference of selection and population growth from patterns of variation in the human genome. Proceedings of the National Academy of Sciences of the United States of America, 102(22):7882–7887.

Wrenbeck, E. E., Azouz, L. R., and Whitehead, T. A. (2017). Single-mutation fitness landscapes for an enzyme on multiple substrates reveal specificity is globally encoded. Nature communications, 8:15695.

Zeng, J., de Vlaming, R., Wu, Y., Robinson, M. R., Lloyd-Jones, L. R., Yengo, L., Yap, C. X., Xue, A., Sidorenko, J., McRae, A. F., Powell, J. E., Montgomery, G. W., Metspalu, A., Esko, T., Gibson, G., Wray, N. R., Visscher, P. M., and Yang, J. (2018). Signatures of negative selection in the genetic architecture of human complex traits. Nature Genetics, 50(5):746–753.

Živković, D., Steinrücken, M., Song, Y. S., and Stephan, W. (2015). Transition Densities and Sample Frequency Spectra of Diffusion Processes with Selection and Variable Population Size. Genetics, 200(2):601–617.

